# Wnt/β-catenin is required for proprioception by coordinating the multi-compartment development of muscle spindles

**DOI:** 10.1101/2024.12.18.629133

**Authors:** Qingyuan Guo, Ava Gatterer, Avital Rosner, Sharon Krief, Fabian S. Passini, Ron Rotkopf, Noa Wigoda, Michael Tsoory, Bavat Bornstein, Elazar Zelzer

## Abstract

Proprioception is essential for the regulation of posture, movement, and musculoskeletal integrity. The muscle spindle, a mechanosensory organ composed of multiple specialized tissues, detects stretch and provides proprioceptive feedback. Despite its importance, the molecular mechanisms that orchestrate the development of the spindle components remain poorly understood.

Here, we reveal the involvement of the Wnt/β-catenin pathway in muscle spindle development. We show that Wnt ligands and their Frizzled receptors are expressed in the developing spindle and that β-catenin is active throughout development in capsule cells and in bag2 intrafusal fibers. Embryonic deletion of β-catenin from capsule and intrafusal fibers induced widespread transcriptomic changes, which led to significant malformations, including impaired capsule cell differentiation, disrupted nuclear organization in intrafusal fibers, and disorganized proprioceptive nerve endings. Postnatal deletion of β-catenin from intrafusal fibers resulted in abnormal nerve endings and impaired proprioceptive function, indicating that the development of proprioceptive afferents is regulated by β-catenin through a non–cell-autonomous mechanism acting in bag2 fibers.

Collectively, our findings position the Wnt/β-catenin pathway as a central regulator that acts through both cell-autonomous and non–cell-autonomous mechanisms to coordinate the development of the various spindle tissues into a functional organ.

## Introduction

The proprioceptive system reports on the relative position and movement of body parts and is therefore essential for control and coordination of movement and posture^1^. In humans, loss of proprioception leads to compromised sensory and motor control resulting in ataxia, abnormal gait, and difficulties in identifying the position of body parts^2^. Furthermore, in recent years, studies in mice show that proprioceptive dysfunction contributes to various musculoskeletal pathologies, including scoliosis, hip dysplasia, and joint contractures^3–6^.

A central component of the mammalian proprioceptive system is the muscle spindle, a small mechanosensory organ that is sparsely embedded within skeletal muscles. The spindle detects changes in muscle length and transmits this information to the central nervous system to establish proprioceptive awareness. A muscle spindle is a complex organ composed of specialized muscle fibers, termed intrafusal fibers, which are innervated by proprioceptive sensory neurons at their central region and by γ-motor neurons at their polar ends. Intrafusal fibers are classified by their nuclear organization and function into three types: bag1, bag2, and chain fibers. The spindle is partially isolated from its surroundings by a capsule rich in extracellular matrix (ECM) that is secreted by capsule cells^7–9^.

Extensive morphological studies have shown that muscle spindle development begins around embryonic day (E) 14, when slow myofibers first contact proprioceptive axons and differentiate into intrafusal fibers in a sequential process^8^. First, nuclear bag fibers are formed, followed by the formation of nuclear chain fibers^10^. In coordination with fiber differentiation, proprioceptive neurons undergo maturation. This morphogenetic process transforms a web-like network into a complex structure termed the annulospiral ending, where neurons wrap around the center of each intrafusal fiber^11^. Capsule development also involves progressive waves of cell differentiation, moving from the center of the spindle to its ends^12^. By contrast, little is known about the molecular aspects of spindle development. Neuregulin 1 (NRG1), expressed by sensory neurons, activates ErbB2 signaling in myofibers to induce the intrafusal genetic program. In return, neurotrophin-3 expressed by intrafusal fibers activates the tropomyosin receptor kinase C (TrkC) receptor on proprioceptive sensory neurons, which is necessary for their survival^13–17^. Recent work has shown that *Lrp4* expression in intrafusal fibers is necessary to maintain the sensory synapses of annulospiral endings and thereby proprioceptive function^18^. Nevertheless, molecular mechanisms underlying the coordinated development of the different spindle tissues are unknown.

Wnt/β-catenin is a central pathway that regulates a variety of cellular processes during development, including cell fate determination, migration, and polarity, as well as tissue homeostasis, regeneration, and pathology. The pathway is activated when Wnt ligands bind to their membrane receptors Frizzled and LRP5/6. This interaction results in the translocation of β-catenin to the nucleus, where it activates transcription of target genes by interacting with TCF/LEF transcription factors^19–20^. Interestingly, Wnt7A expression was detected in γ-motor neurons of the muscle spindle, whereas Frizzled5 was expressed in the capsule^21^. These observations suggest that Wnt/β-catenin signaling may be involved in spindle development.

In this study, we investigate the role of the Wnt/β-catenin pathway in the development of muscle spindle tissues. We first identify expression of both Wnt ligands and Frizzled receptors in distinct spindle compartments and show that β-catenin remains active in capsule cells and bag2 fibers throughout development. We then use several mouse models to delete β-catenin from specific spindle tissues either pre- or postnatally. These perturbations led to extensive transcriptomic alterations, as well as to both cell-autonomous and non-autonomous effects on capsule cell differentiation, nuclear organization in intrafusal fibers, and morphogenesis of proprioceptive nerve endings. These findings establish β-catenin as a central regulator of muscle spindle development and uncover a set of previously unrecognized genes involved in this process.

## Results

### The Wnt/ β-catenin pathway is active in muscle spindles throughout development

To identify molecular pathways that may regulate muscle spindle development, we performed pathway analysis using Ingenuity Pathway Analysis (IPA)^22^ to RNA-seq data we had recently obtained from isolated spindles^12^. We identified numerous upstream regulators of genes enriched in spindles, including the top candidates *Pou4f1, Dnmt3b,* and *Ctnnb1* (Table S1). Of note, *Ctnnb1* encodes β-catenin, which is a central intracellular signal transducer in the Wnt signaling pathway^19–20^. Indeed, IPA revealed enrichment of multiple components of the Wnt/β-catenin pathway (Figure 1A), including Wnt ligands *Wnt1, Wnt2b, Wnt3, Wnt6, Wnt7b and Wnt10a, Wnt10b, Wnt16*, Frizzled receptors *Fzd1, Fzd2, Fzd3, Fzd5, Fzd9* and *Fzd10*, and TCF/LEF transcription factors (Figure 1B and Table S2). Based on these results, we hypothesized that Wnt/β-catenin signaling plays an important role in muscle spindle development.

**Figure 1.**
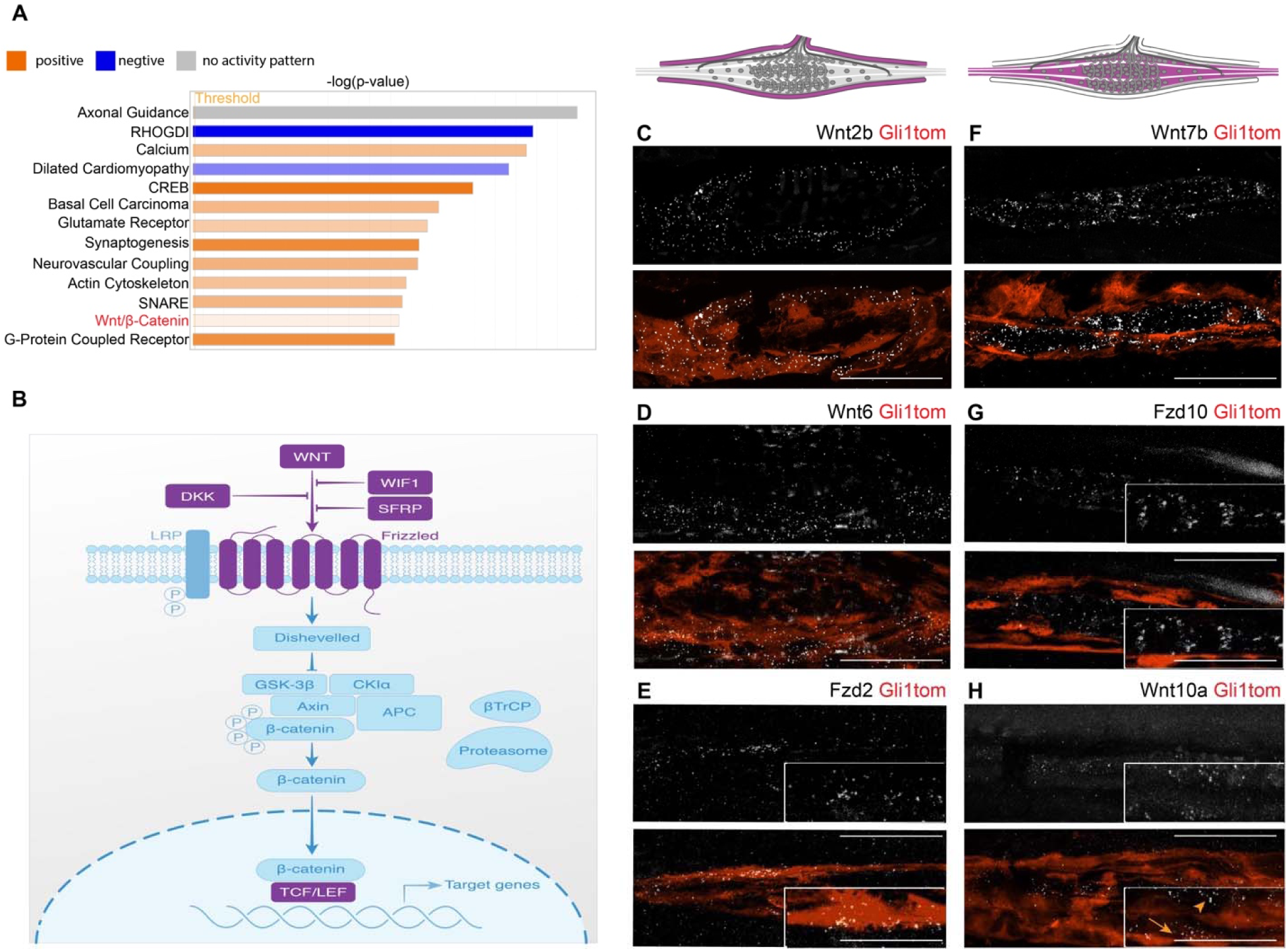
Transcriptomic and HCR analyses reveal expression of Wnt/β-catenin pathway components in muscle spindles. (A) Ingenuity Pathway Analysis of differentially expressed genes (DEGs) between muscle spindle and muscle samples. (B) Schematic representation of Wnt/β-catenin pathway. Purple blocks represent genes upregulated in spindles compared to muscles. (C-H) Confocal images of longitudinal sections of deep masseter muscles (C-D, F, H) and whole-mount of EDL muscles (E, G) from P33 *Gli1-CreER^T2^,Rosa26-tdTomato* mice. Expression of the indicated Wnt/β-catenin pathway genes was detected by HCR probes following tamoxifen administration at P26. *Wnt2b*, *Wnt6* and *Fzd2* are expressed in capsule cells (C-E; inset in E shows enlargement of *Fzd2* expression). *Wnt7b* and *Fzd10* are expressed in intrafusal fibers (F,G; inset in G shows enlargement of *Fzd2*), and *Wnt10a* is expressed in both (H; inset in G shows enlarged image). Arrows points at capsule expression and arrowheads point at intrafusal expression. Top: schematic representation of muscle spindles showing in purple the tissue in which the indicated genes were expressed. Scale bars: 50 µm; 100 µm for enlarged images.

To validate our transcriptomic results and to determine the spatial distribution of some of these molecules, we applied single-molecule in situ hybridization chain reaction (HCR) to frozen sections from deep masseter muscles and whole-mount extensor digitorum longus (EDL) muscles. To outline the various spindle tissues, we used *Gli1-CreER^T2^;tdTomato* to mark capsule cells^12^ (Figure 1C-H) and *Calb1-Cre;tdTomato* to mark intrafusal fibers^23^ (Figure S1). Results showed that *Wnt2b*, *Wnt6,* and *Fzd2* were expressed by capsule cells (Figure 1C-E), whereas *Wnt7b* and *Fzd10* were expressed by intrafusal fiber cells (Figure 1F-G and Figure S1A,C). *Wnt10a* was expressed in both capsule cells and intrafusal fibers (Figure 1H and Figure S1B). The variable expression of Wnt and Fzd molecules in different spindle tissues suggests that Wnt/β-catenin signaling has multiple roles in muscle spindle development.

Next, to validate Wnt/β-catenin pathway activity in the developing spindle, we used *Axin2*-membrane GFP (mGFP) reporter mice. *Axin2* is a direct target of Wnt/β-catenin and *Axin2-mGFP* mice report pathway activity faithfully^24^. As seen in Figure 2A, we detected GFP signal in both extrafusal fibers and muscle spindles at postnatal day (P) 0 and P5. At P25 and P40, the GFP signal was lost in extrafusal fibers but maintained in muscle spindles. To determine in which spindle tissue Wnt signaling was active, we stained EDL muscles from *Axin2-mGFP* mice using antibodies against MyHC7B and GLUT1, markers for bag2 fiber and capsule cells, respectively^12, 25^. As shown in Figure 2B,C and Figure S2, from P5 and onwards, Wnt/β-catenin activity was specific to bag2 fibers and capsule cells. Overall, these results show ongoing Wnt signaling in several spindle tissue types, further supporting diverse roles in spindle development.

**Figure 2.**
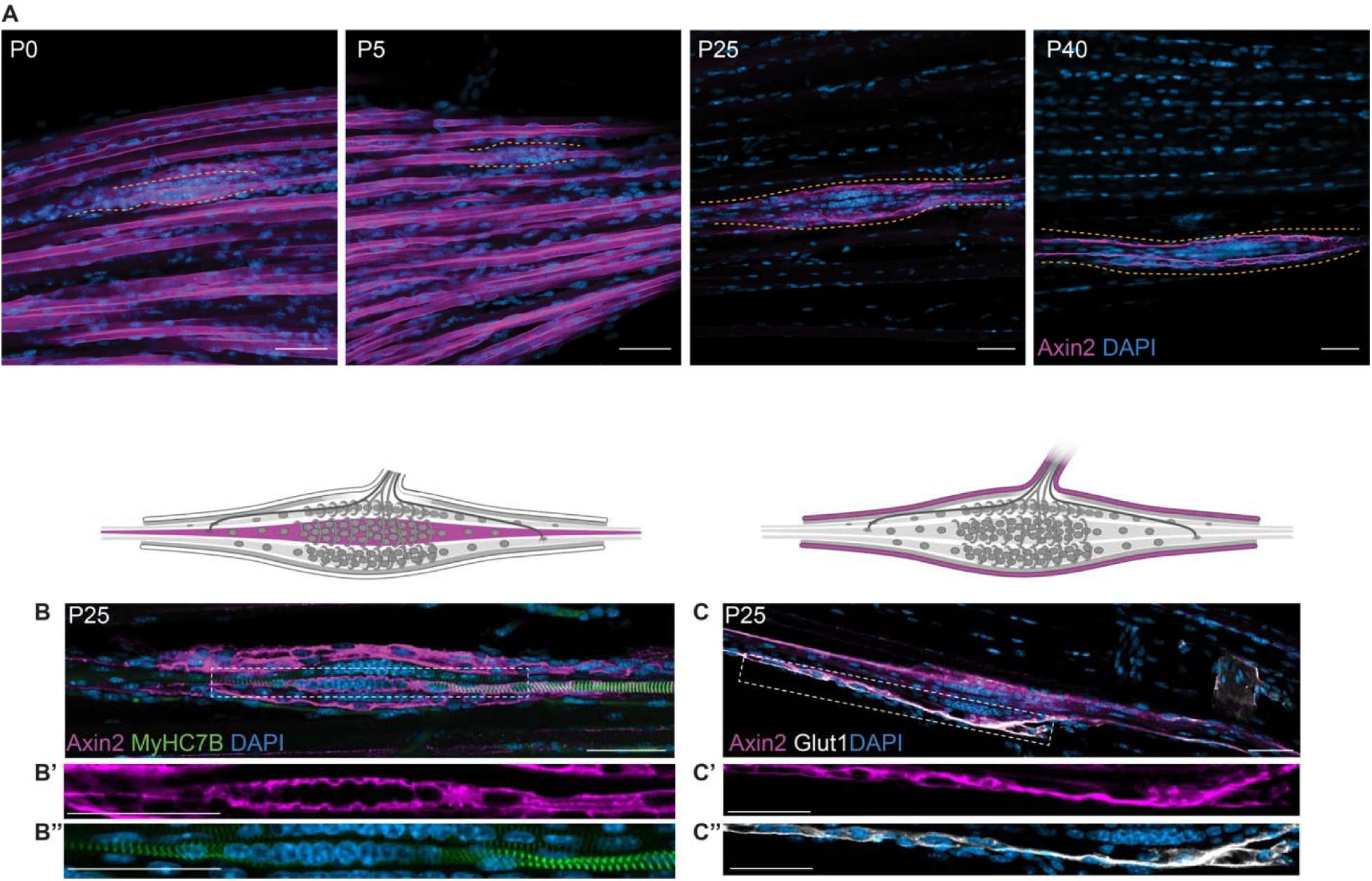
β-catenin activity is detected in muscle spindles throughout development. (A) Confocal images of whole-mount EDL muscles from P0, P5, P25 and P40 *Axin2-mGFP* mice stained with antibodies against GFP (magenta) and DAPI (blue). Dotted lines demarcate equatorial region of spindles. Scale bars: 50 µm. (B-C’’) Confocal images of whole-mount EDL muscles from P25 *Axin2-mGFP* mice stained with antibodies against GFP, DAPI and MyHC7B (green, B-B’’) or GLUT1 (white, C-C’’). Above are schematic representations of muscle spindles with the tissue analyzed in purple. Scale bars: 50 µm.

### Deletion of β-catenin from capsule and intrafusal fibers results in transcriptomic changes in the capsule, intrafusal fibers and proprioceptive nerve endings

Our findings that Wnt ligands and their Frizzled receptors are expressed in the developing spindle and that β-catenin is active in both capsule and intrafusal fibers throughout postnatal development prompted us to evaluate the transcriptional program regulated by β-catenin by analyzing β-catenin–mutant spindles. For this purpose, we crossed β*-catenin^loxP/loxP^*mice to *Egr3-Cre* mice, thereby deleting β-catenin from both intrafusal fibers and capsule cells^26,27^. Given that *Egr3-Cre* also targets β-catenin in the periphery of the spindle, including the interstitial cells of the extrafusal fibers (Figure S3), we conducted comparative RNA-seq of masseter muscle spindles and surrounding extrafusal fibers from *Egr3-Cre*, β*-catenin^loxP/+^* control mice versus *Egr3-Cre,* β*-catenin^loxP/loxP^*mutant mice. Principal component analysis (PCA) showed a clear separation between control and mutant spindles (Figure S4). Differential gene expression analysis identified 747 downregulated and 297 upregulated genes in mutant spindles (Fig 3A).

**Figure 3.**
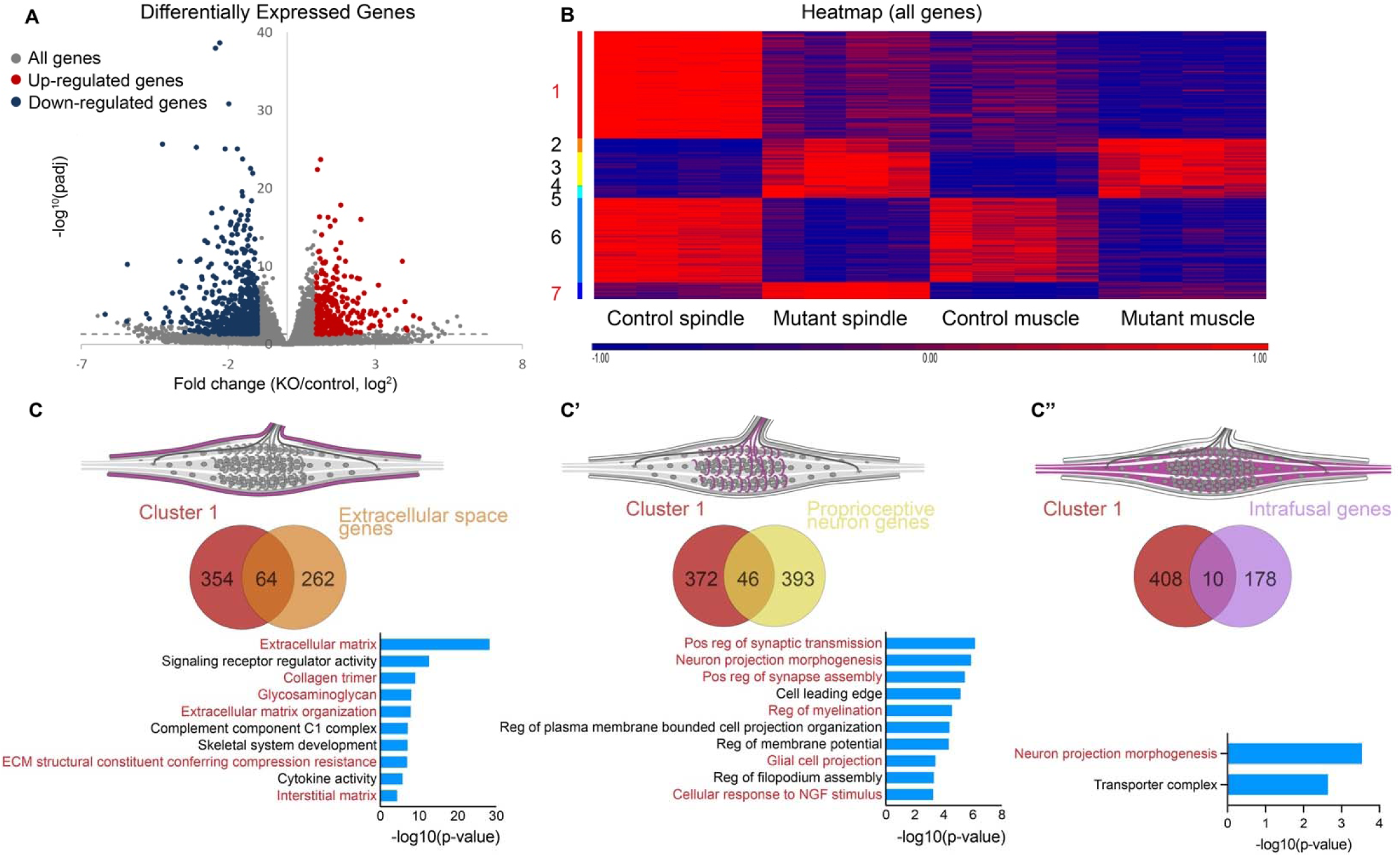
Transcriptomic analysis of *Egr3-*β*-catenin* mutant and control tissues. (A) Volcano plot of DEGs between mutant *Egr3-Cre,* β*-catenin^loxP/loxP^*and control *Egr3-Cre,* β*-catenin^loxP/+^*spindles. (B) Heatmap of DEGs between control and mutant spindle and muscle tissues. (C-C’’) Top: schematic representation of the muscle with the analyzed tissue in purple. Middle: A Venn diagram showing overlapping genes between differentially downregulated genes in mutant spindles and extracellular space genes (C), proprioceptive neurons (C’) and intrafusal genes (C’’). Bottom: GO analysis for enriched biological processes in the overlapping datasets.

To identify spindle-specific differentially expressed genes (DEGs), we performed K-means clustering for spindle and extrafusal fiber samples. As seen in Figure 3B, cluster 1 contained 418 spindle-specific genes that were downregulated and cluster 7 contained 65 spindle-specific genes that were upregulated in mutant mice, as compared to the control (Table S3). IPA of cluster 1 showed enriched pathways associated with connective tissue, neuronal and synaptic pathways (Table S4). Interestingly, among the neuronal and synaptic pathways, IPA identified genes encoding ligand and receptor pairs crucial to both synaptogenesis and axon guidance, including the NGFR-NGF, Eph-ephrin, neurexin-neuroligin, and semaphorin-plexin signaling pathways.

Next, to associate these changes with the various spindle tissue types, we cross-referenced cluster 1 with published tissue type-specific transcriptomic data^11^. We identified 64 extracellular, 46 proprioceptive and 10 intrafusal genes (Fig 3C-C’’, Table S5). Gene ontology (GO) analysis of capsule-associated extracellular genes showed terms like “extracellular matrix” and “collagen trimer”, including genes such as *Loxl4*, *Vcan*, *Fbln1*, *Col9a2*, *Col12a1*, *Col4a5*, *Col18a1* and *Col8a1*. GO terms for proprioceptive neuron genes were “positive regulation of synaptic transmission” and “neuron projection morphogenesis”, including genes like *Ntrk3* and *Ngfr*. Surprisingly, for intrafusal genes, the most enriched GO term was “neuron projection morphogenesis”, including genes such as *Ngfr* and *Wnt7b*, suggesting intrafusal regulation of neuronal tissue development.

Further analysis of cluster 1 uncovered a set of genes previously unknown to be involved in spindle biology. This list contains gap junction genes, including *Gja5*, *Gjb5*, *Gjb1*, *Gjb6* and *Gjb2*; the tight junction gene *Ocln* and the associated *Cgnl1*; retinol transporter gene *Stra6*; nuclear movement-associated genes such as *Pard6g* and *Lmnb2*.

Collectively, these results show that the absence of β-catenin from capsule and intrafusal fibers led to transcriptomic changes in ECM, connective tissue and neuronal genes.

### Deletion of β-catenin from capsule and intrafusal fibers results in cell-autonomous and cell non-autonomous phenotypes in muscle spindles

To understand the consequences of the transcriptomic changes in β-catenin mutated spindles, we studied the morphological and molecular effects of β-catenin loss on the development of various spindle tissues in P25 mice, in which muscle spindles morphogenesis is largely complete^8,12^. First, we examined the protein expression of GLUT1 and VCAN, markers for early and advanced capsule differentiation, respectively^12^. As shown in Figures 4A and 4E, GLUT1 was expressed throughout control muscle spindles, whereas in mutant spindles its expression was restricted to the equatorial region. Moreover, unlike in controls, VCAN expression was reduced in mutant spindles (Figure 4B), suggesting that capsule cells failed to differentiate. Consistent with the observed reduction in ECM, mutant spindles appeared thinner than controls (Figures 4A and 4E’).

**Figure 4.**
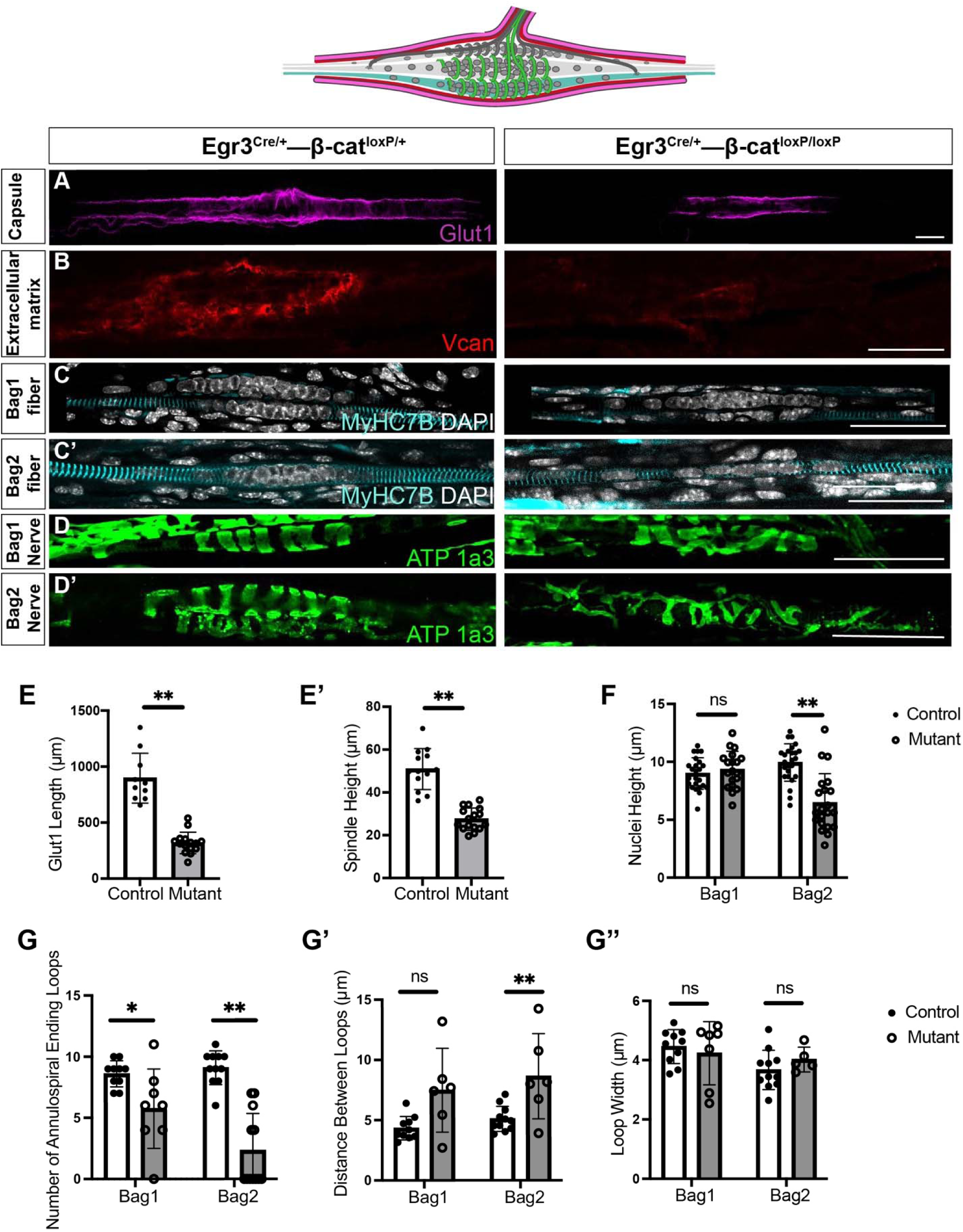
Deletion of β-catenin from capsule and intrafusal fibers results in morphological abnormalities in three spindle tissues. (A-D’) Confocal images of P25 muscle spindles from mice of *Egr3-Cre,* β*-catenin^loxP/loxP^* and control *Egr3-Cre,* β*-catenin^loxP/+^*mice. Capsules were stained with anti-GLUT1 antibodies in whole-mount gluteus muscles, ECM was stained with anti-VCAN antibodies in longitudinal sections from deep masseter muscles, bag2 intrafusal fibers were stained with anti-MyHC7B antibodies in whole-mount EDL muscles, and proprioceptive neurons were stained with anti-ATP1a3 antibodies in whole-mount gluteus muscles. Scale bars: 50 µm. (E-G’’) Graphs showing quantitative analyses of control and β-catenin mutant muscle spindle morphology. (E): GLUT1 expression was restricted to the equatorial regions in mutants (**P<0.01, n=3 mice, 10 control spindles, 15 mutant spindles). Gluteus muscles were used for quantification. (E’): Decreased distance between GLUT1^+^ capsule layers indicates a shorter spindle height in mutants (**P<0.01, n=3). Gluteus muscles were used for quantification. (F): Abnormal nuclear aggregation and shorter nucleus height in the equatorial regions of mutant intrafusal fibers (**P<0.01, n=4 mice, 25 control spindles, 21 mutant spindles). EDL muscles were used for quantification. (G-G’’) Although the width of annulospiral endings was not affected, the distance between loops and the number of loops per spindle was decreased in mutants (**P<0.01, *P<0.05, n=3 mice, 11 control spindles, 13 mutant spindles). EDL muscles were used for the quantification.

Next, we studied intrafusal fiber development. In agreement with the activation patterns of β-catenin, we found abnormal nuclear aggregation in the equatorial region of mutant bag2 fibers, whereas in bag1 fibers nuclear aggregation was similar to wild type (Figure 4C, C’, F). These results suggest that β-catenin is necessary for bag2 fiber development.

Finally, we studied the development of the annulospiral endings of type Ia proprioceptive neurons. As seen in Figure 4D-D’, staining with antibody against ATP1a3, a marker for proprioceptive endings, revealed stereotypical annulospiral loops in control proprioceptive neurons, whereas β-catenin mutant spindles exhibited abnormal annulospiral morphology (Figure 4G-G’). While mutant loops had similar widths, loop number was reduced and the distance between loops increased (Figure 3G-G’’). Interestingly, although these morphological alterations were seen in annulospiral endings of both bag1 and bag2 fibers, they were more severe in the latter.

Overall, our findings indicate that Wnt/ β-catenin activity is necessary for the development of the capsule, intrafusal fibers, and proprioceptive neuron, suggesting that β-catenin regulates muscle spindle development both autonomously and non-autonomously.

### Intrafusal fiber-specific deletion of β-catenin leads to abnormal annulospiral endings

To detect the tissue from which β-catenin exerts its cell non-autonomous effect on nerve endings, we used the *HSA^CreER^* line^28^, which allows temporal targeting of extrafusal and intrafusal fibers. Previous studies using this strain to delete β-catenin postnatally have not identified a clear phenotype in extrafusal fibers^29^. We therefore expected that any spindle phenotype would represent a direct role of β-catenin in intrafusal fibers.

For this experiment, we crossed *HSA^CreER/+^,* β*-catenin^loxP/+^* mice with β*-catenin^loxP/loxP^*mice and administered tamoxifen at P5 before collecting spindles at P25. As shown in Figure 5A-B, expression patterns and levels of both capsule markers GLUT1 and VCAN were unaffected in β-catenin mutant mice. Similarly, the distance between GLUT1^+^ layers was normal, indicating that the capsule of *HSA^CreER^-*β*-catenin* mutant spindles develops normally (Fig. 5E-E’). Comparison of intrafusal fibers between control and mutant spindles showed no difference in nuclear aggregation (Fig. 5C-C’, F). Interestingly, however, mutant annulospiral endings of bag2 fibers were abnormal, with fewer and more spaced loops, whereas annulospiral endings of bag1 fibers were intact (Fig. 5D-D’, G-G’’). These results suggest that the development of proprioceptive nerve endings is regulated by β-catenin through a non–cell-autonomous mechanism acting in bag2 intrafusal fibers.

**Figure 5.**
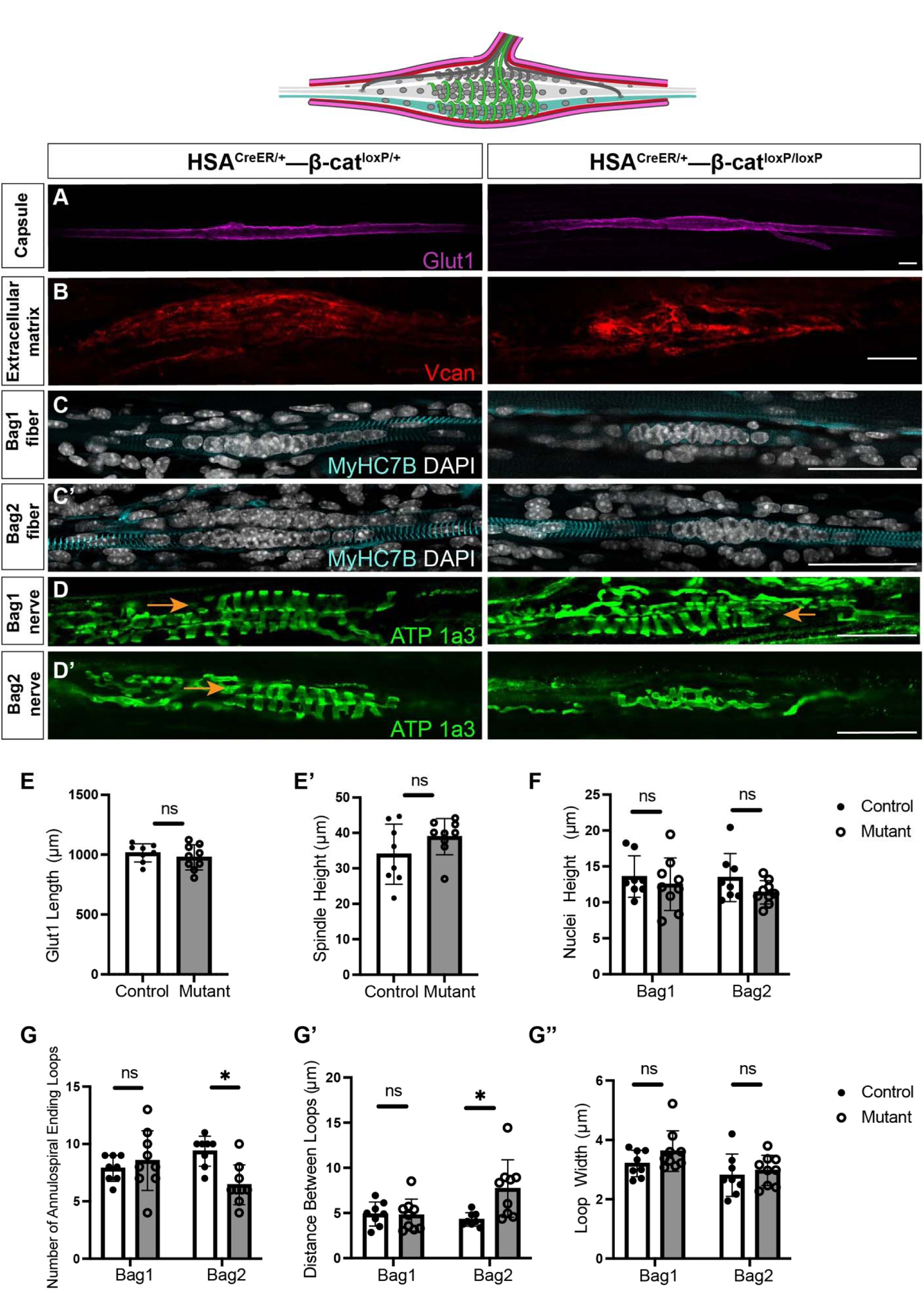
Deletion of β-catenin from intrafusal fibers results in disorganized annulospiral endings. (A-D’) Confocal images of muscle spindles from *HSA^CreER^-*β*-catenin* mutant and control mice. Capsules were stained with anti-GLUT1 antibodies in whole-mount EDL muscles, ECM was stained with anti-VCAN antibodies in longitudinal sections from gluteus muscles, bag2 intrafusal fibers were stained with anti-MyHC7B antibodies in whole-mount EDL muscles, and proprioceptive neurons were stained with ATP1a3 antibodies in whole-mount EDL muscles. Arrows point at bag1 fiber in (D) and bag2 fiber in (D’). Scale bars: 50 µm. (E-G’’) Graphs showing quantitative analyses of control and β-catenin mutant muscle spindle morphology. (E-F): Expression of GLUT1, spindle height as measured by the distance between GLUT1^+^ capsule layers, and nuclear aggregation in intrafusal fibers were not significantly different between controls and mutants (n=3 mice, 8-9 spindles). (G-G’’): Although the width of annulospiral endings was similar, the distance between loops and the number of loops per spindle was decreased in mutants (*P<0.05, n=3 mice, 8-9 spindles).

### Intrafusal fiber-specific deletion of β-catenin leads to impaired proprioceptive function

Given the malformation of proprioceptive nerve endings in *HSA^CreER/+^,* β*-catenin^loxP/loxP^* mice, we next investigated proprioceptive function by using the CatWalk system. Analyzed coordination-associated parameters included step sequence, number of patterns, regularity index, print position, stride length, support and base of support (BOS)^30–31^. Step sequence analysis showed that in comparison with controls, mutant mice used less the Ca walking pattern, which requires more contralateral coordination, and more Ab patterns, which relies more on ipsilateral coordination, suggesting impaired interlimb coordination (Fig. 6A, S5A). In addition, as compared to controls, while walking, mutant mice supported themselves more on all four paws and less on a single paw, reflecting a need for more support (Fig. 6D, S5B). Similarly, BOS analysis indicated a trend for a wider front paw distance in β*-catenin* mutants, further suggesting that mutants seek more support while walking (Fig. 6F). No differences were found between genotypes in the number of patterns used, regularity index, stride length or print position (Fig. 6B-C,E,G). Notably, no difference in mean walking speed was observed between mutants and controls, suggesting that the deletion of β-catenin did not result in a global mobile dysfunction, but rather in defects in coordination and balance. Overall, theses data suggest that the cell non-autonomous effect of β-catenin on proprioceptive nerves results in impaired proprioceptive function.

**Figure 6.**
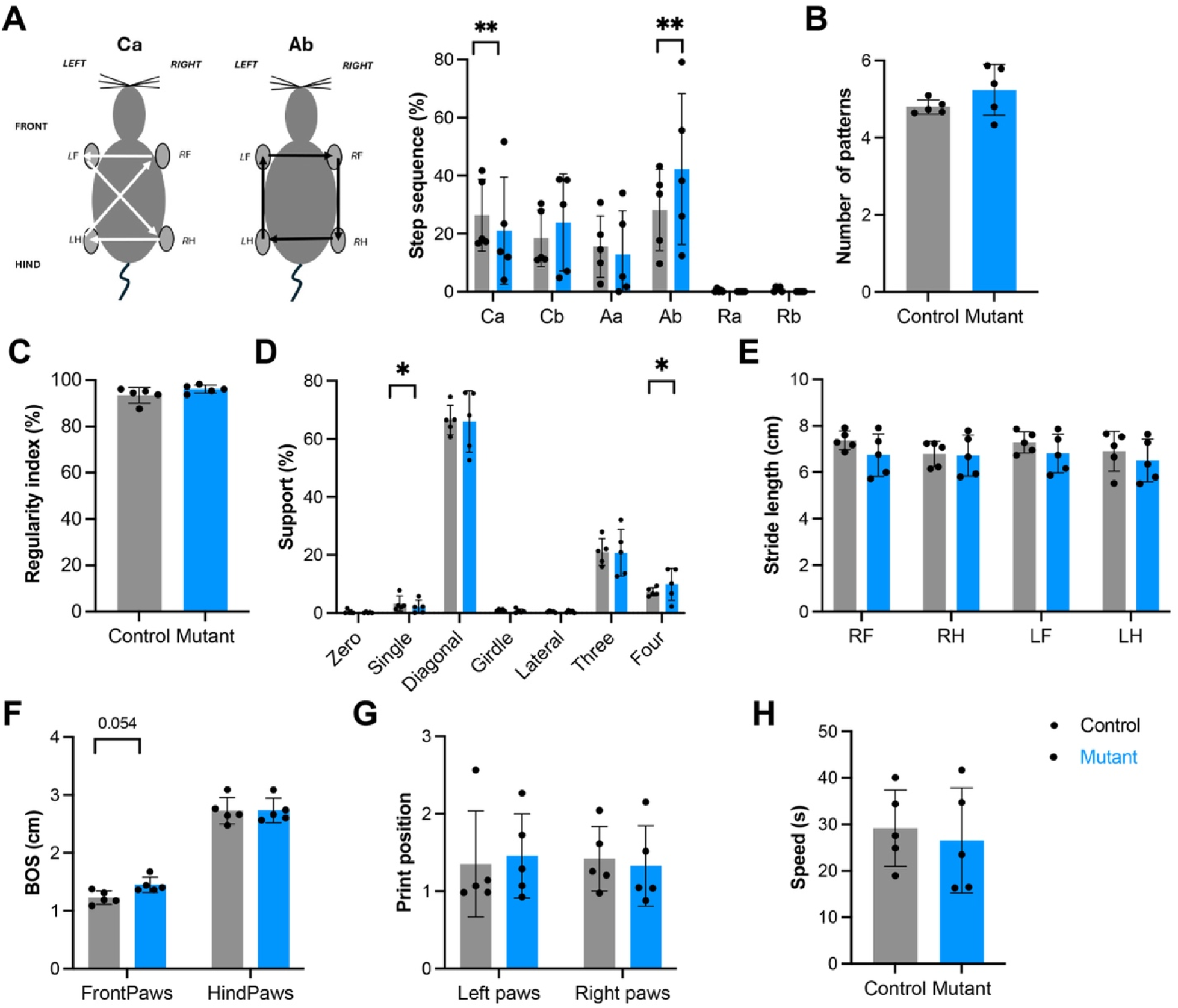
Deletion of β-catenin from intrafusal fibers led to impaired proprioceptive function. Stride and gait were assessed using the CatWalk system. (A,B) Step sequence pattern is the sequence of paw placements and includes six patterns: Aa, alternative a; Ab, alternative a; Ca, cruciate a; Cb, cruciate b; Ra, rotary a; Rb, rotary b. (C) Regularity index defines the percentage of steps that are part of a recognized sequence pattern out total number of paw placements. (D) Support is the number and position of paws supporting the animal in each step. (E) Stride length is the distance between consecutive placements of the same paw. (F) Base of support (BOS) represents the distance between either front or hind paws. (G) Print position is the distance between consecutive placements of ipsilateral hind and front paws. (H) The mean walking speed. *HSA-*β*-catenin* mutants used the Ca step sequence pattern less frequently and the Ab step sequence pattern more frequently (Ca, p=0.002; Cb, p=0.174; Aa, p=0.456; Ab, p=0.003; Ra, p=0.937; Rb, p=0.783). The mutants also needed more support (zero, p=0.666; single, p=0.038; diagonal: p=0.924; girdle, p=0.200; lateral, p=0.787; three, p=0.773; four, p=0.031). No difference was found between *HSA-*β*-catenin* mutants and controls in the number of patterns (p=0.256), regularity index (p=0.176), stride length (RF, p=0.154; RH, p=0.880; LF, p=0.267; LH, p=0.368) BOS (front paws, p=0.054; hind paws, 0.951), print position (left paws, p=0.775; right paws, p=0.799) or mean speed (p=0.579). n=5 in each group; each dot represents a mouse; data are represented as the mean ± SD.

## Discussion

Although the muscle spindles are at the core of the proprioceptive system, surprisingly little is known about the molecular mechanisms underlying their development. Here, we show that Wnt/β-catenin signaling is active in developing spindles and is required for the proper formation of specific components of the spindle. Deletion of β-catenin from the embryonic capsule and intrafusal fibers led to extensive transcriptomic changes, which were accompanied by pronounced malformations in the capsule, intrafusal fibers, and proprioceptive nerve endings. Furthermore, loss of β-catenin specifically in bag2 intrafusal fibers resulted in abnormal development of proprioceptive nerve endings and impaired proprioceptive function, indicating a non–cell-autonomous effect. Collectively, these findings establish both cell-autonomous and non–cell-autonomous roles for β-catenin in muscle spindle development.

Our analysis of RNA-seq data revealed enrichment of multiple components of the Wnt/β-catenin pathway in the muscle spindle. Consistently, we observed diverse expression patterns of numerous Wnt ligands and Frizzled receptors within the spindle. These findings support a central role for this pathway in muscle spindle development and raise the possibility that different Wnt– Fzd combinations mediate distinct, tissue-specific functions. For example, *Wnt7b*, *Wnt10a*, and *Fzd10* were all expressed in the spindle capsule. Given that WNT7B/WNT10A have been shown to bind directly to FZD10^32–33^, a dedicated signaling axis may guide capsule development. Indeed, we found that β-catenin–mutant capsule cells fail to differentiate and to secrete ECM, likely impairing capsule structure and function.

Previous studies suggest that the muscle spindle capsule functions as a selective diffusion barrier, similar to the blood–brain barrier (BBB) and blood–nerve barrier (BNB)^34–36^. Structurally, the capsule mirrors key features of these classical barriers, being enriched in gap junctions, tight junctions, and ECM components^37^. Our transcriptomic analysis further identified reduced expression of *Stra6*, a retinol transporter, and *Glut1*, a glucose transporter, suggesting that these transporters may regulate nutrient flux across the spindle capsule, similar to their established roles in the BBB^38–40^. In addition, we observed decreased expression of multiple gap junction, tight junction, and ECM components. Notably, several of these genes, such as *Ocln* and *Cgnl*, are essential for the structural integrity and permeability of the BBB^41–42^, raising the possibility that they play analogous roles in maintaining the selective permeability and structural integrity of the muscle spindle capsule.

One particularly interesting observation from our study is the selective activity and function of β-catenin in bag2, but not in bag1 intrafusal fibers. Bag2 fibers are the first intrafusal fibers to form during spindle development, with bag1 fibers emerging later^10^. Physiologically, bag2 fibers mediate static sensitivity by encoding muscle length, whereas bag1 fibers mediate dynamic sensitivity, responding to the velocity of stretch. These distinct physiological roles arise from a collection of molecular and structural differences between the two fiber types^43–44^. Despite these pronounced differences, little is known about the molecular mechanisms that specifically regulate each intrafusal fiber subtype. Our finding that β-catenin is selectively active in bag2 fibers, where it is required for both nuclear aggregation and the proper establishment of proprioceptive nerve endings, provides the first bag2 fiber-specific regulatory mechanism and indicates that the two fiber types are indeed governed by distinct pathways.

While nuclear aggregation is a hallmark of intrafusal fibers, little is known about the mechanisms that regulate this process. The downregulation of *Pard6g*, a polarity factor from the Par6 family that interacts with the dynein/dynactin complex at the nuclear envelope, and *Lmnb2*, which encodes a B-type nuclear lamin required for nuclear motion, in β-catenin–mutant spindles offers a potential mechanistic explanation for the observed loss of nuclear aggregation in bag2 fibers^45–46^.

Cross-talk between intrafusal fibers and proprioceptive neurons has been previously described. For example, intrafusal neurotrophin-3 (NT-3) is required for the survival of proprioceptive neurons through activation of the TrkC receptor during embryonic development^17^. Similarly, the expression of LRP4 in intrafusal fibers has been shown to be essential for the development, maintenance, and aging of proprioceptive sensory synapse morphology and function^18^. Our finding that β-catenin activity in bag2 fibers regulates sensory endings adds another key component to this bidirectional communication between intrafusal fibers and proprioceptive afferents.

Regarding the potential mechanisms underlying this cross-talk, our transcriptomic analysis of β-catenin mutant spindles highlights several candidate molecules and pathways. We observed reduced expression of genes involved in axon guidance and synaptogenesis, including components of the ephrin–Eph, semaphorin–plexin, and neurexin–neuroligin pathways^47–49^. The transcription of two downregulated guidance genes, *EphB2* and neuroligin 3, is regulated by Wnt/β-catenin signaling in other biological systems, including the intestinal epithelium and hippocampus^50^. Together, these findings suggest that β-catenin influences proprioceptive nerve endings through multiple interconnected pathways, revealing new molecular mechanisms by which intrafusal fibers shape spindle innervation.

In summary, we establish Wnt/β-catenin signaling as a key regulator of the coordinated development of the various components of the muscle spindle. In particular, we demonstrate its essential role in capsule differentiation, as well as its specific requirement in bag2 fibers for proper nuclear aggregation and for the formation and organization of proprioceptive nerve endings around them. Finally, by integrating transcriptomic profiling with the distinct morphological phenotypes observed in β-catenin mutants, we uncovered novel genes enriched in muscle spindles and inferred their potential functions. Collectively, these findings offer multiple molecular entry points into the development of individual spindle tissues, including capsule, intrafusal fibers, and proprioceptive synapses, and provide insight into the mechanisms mediating the communication between these tissues.

## Methods

### Mouse lines

The generation of *Axin2-mGFP* (The Jackson Laboratory, #037313)^24^, *calbindin1-Cre*^23^, *Egr3-Cre*^26^, *HSA-MCM* (also known as Tg(ACTA1-cre/Esr1*)2Kesr/J; The Jackson Laboratory, #025750)^28^, *Gli1-CreER^T2^* (The Jackson Laboratory, #007913)^12^, *Rosa26-tdTomato* (The Jackson Laboratory, #007909)^51^ and β*-catenin^flox^* (aka B6.129-*Ctnnb1^tm2Kem^*/KnwJ; The Jackson Laboratory, #004152)^27^ has been described previously. For HCR of Wnt genes, *calbindin1-Cre* mice and *Gli1-CreER ^T2^* mice were crossed with *Rosa26-tdTomato* mice. To induce Cre-mediated recombination in *Gli1-CreER^T2^:Rosa26-tdTomato* mice, 200 mg/kg of 50 mg/mL tamoxifen (Sigma-Aldrich, T5648) dissolved in corn oil (Sigma-Aldrich, C-8267) was administered at indicated age via oral gavage (Fine Science Tools). To generate *Egr3* control and mutant mice, *Egr3-Cre* mice were crossed with β*-catenin^flox/flox^, Rosa26-tdTomato* mice. To generate HSA control and mutant mice, *HSA-MCM* mice were crossed with β*-catenin^flox/flox^, Rosa26-tdTomato* mice. To induce Cre-mediated recombination, 7 µL of 50 mg/mL tamoxifen in corn oil was administered by oral gavage for 3 consecutive days starting from P5. All animals were sacrificed by CO_2_ inhalation, fixed overnight shaking in 4% paraformaldehyde (PFA)/PBS at 4°C, washed in PBS and stored at 4°C in PBS.

Mice were given free access to water and standard mouse chow with 12-h light/dark cycles. All animal procedures were approved by the Institutional Animal Care and Use Committee of Weizmann Institute. Ear genomic DNA was used for genotyping by PCR.

### Immunofluorescence

For cryosection immunofluorescence, tissues were processed as previously described^12^. In brief, fixed deep masseter muscles and gluteus muscles were dissected and placed in 30% sucrose overnight with shaking. The muscles were then embedded in OCT and sectioned at 30 µm thickness using a cryostat. Cryosections were dried, post-fixed in 4% PFA, permeabilized with PBS containing 0.3% Triton X-100, washed with PBS containing 0.1% Tween-20 (PBST) and blocked with 7% goat/horse serum and 1% bovine serum albumin (BSA) in PBST. The sections were then incubated with primary antibodies (Table S6) overnight at 4°C, followed by a 1-hour incubation with secondary antibodies conjugated with fluorophores (Table S6) at room temperature. The sections were counterstained with DAPI and mounted with Immuno-mount aqueous-based mounting medium (Thermo Fisher Scientific). Imaging was performed with LSM 800 or LSM900 confocal microscope (Zeiss).

For whole-mount immunofluorescence, muscles were subjected to an optical tissue clearing protocol as described previously^11^. Shortly, fixed EDL muscles were dissected and embedded into an A4P0 hydrogel (4% acrylamide, 0.25% 2’-azobis[2-(2-imidazolin-2-yl)propane]dihydrochloride in PBS) with shaking at 4°C overnight. Then, samples were polymerized in hydrogel for 3 h at 37°C and the hydrogel was removed by washing in PBS. Next, samples were transferred to 10% SDS (pH 8.0) with 0.01% sodium azide, shaking gently at 37°C for 2 days. Cleared samples were then washed in wash buffer, blocked with CAS block for 5 h at 37°C with shaking, and incubated with primary antibodies (Table S6) for 4 days at 37°C with shaking. They were then incubated with secondary antibodies (Table S6) and DAPI (8 µg/ml) for 2 days at 37°C with shaking. For clearing and mounting, samples were incubated in 500 µl of refractive index matching solution (RIMS; 74% wt/vol Histodenz in 0.02 M phosphate buffer) for 1 day at room temperature with gentle shaking. Samples were then embedded in RIMS and imaged using a Zeiss LSM800 or LSM900 confocal microscope. Images were processed with ImageJ 1.51 (National Institute of Health).

### Single-molecule *in situ* hybridization chain reaction (HCR)

For cryosection HCR, deep masseter muscles were dissected, transferred to 30% sucrose overnight with shaking, then embedded in OCT and sectioned at a thickness of 30 µm using cryostat. Cryosections were processed for HCR smFISH as described in http://www.molecularinstruments.com. Briefly, slides were incubated with the specified probes overnight, washed and incubated with fluorophore-conjugated amplifiers. The sections were counterstained with DAPI, mounted with Immuno-mount, and imaged using a LSM 800 or LSM900 confocal microscope (Zeiss).

For whole-mount HCR, EDL muscles were dissected, subjected to optical tissue clearing, and processed for HCR smFISH as previously described^52^. Shortly, samples were incubated in probe solution overnight at 37°C, washed, and then incubated with fluorescent amplifiers overnight at room temperature. Next, samples were washed and counterstained with DAPI (8 µg/ml) for 2 h at room temperature. Finally, samples were moved to 500 µl RIMS overnight at room temperature and embedded in RIMS for imaging. Images were captured by LSM800 or LSM900 confocal microscope (Zeiss).

### Bulk RNA sequencing

For RNA-seq analysis, tissues were processed as described before^12^. Deep masseter muscles, each containing about 20 spindles, were dissected from *Egr3-Cre;TdTom;catenin* control and mutant mice. Then, bundles of spindles and adjacent extrafusal muscle fibers were micro-dissected and frozen on dry ice. For each sample, we pooled the same tissue from two animals of the same genotype. Each genotype had four replicates for each tissue, four samples of spindles and four samples of extrafusal fibers for both mutants and controls.

#### Muscle spindle isolation

To isolate muscle spindles from deep masseter muscles, spindles were dissected in ice-cold Liley’s solution (<u>Liley, 1956</u>) (NaHCO_3_ 1 g, KCl 0.3 g, KH_2_PO_4_ 0.13 g, NaCl 0.2 g, CaCl_2_ 1 M 2 ml in total of 1 L DDW).

#### Sample preparation

Total RNA was extracted using TRIzol Reagent followed by chloroform phase separation and processing with the RNeasy Micro kit (Qiagen). The quality and concentration of RNA were assessed using NanoDrop and TapeStation. RNA-seq libraries were generated at the Crown Genomics Institute of the Nancy and Stephen Grand Israel National Center for Personalized Medicine, Weizmann Institute of Science, following the INCPM-mRNA-seq protocol. The libraries were quantified by Qubit and TapeStation. Sequencing was conducted on an Illumina NextSeq instrument with a single end 75 cycles high output kit, yielding approximately ∼20 M reads or more per sample.

#### Analysis of RNA-seq data

Transcriptomic data from four replicates of spindle samples and four muscle samples from mutants and controls were analyzed. A user-friendly Transcriptome Analysis Pipeline (UTAP) version 1.10.1 was used for analysis^53^. Reads were mapped to the *M. musculus* genome (GRCm38), using STAR (v2.5.2b)^54^ and RefSeq annotation. Only reads with unique mapping were considered for further analysis. Gene expression was calculated and normalized using DESeq2 version 1.16.1^55^, using only genes with a minimum of five reads in at least one sample. Raw p-values were adjusted for multiple testing^56^. A gene was considered differentially expressed if it passed the following thresholds: minimum mean normalized expression of 5, adjusted p-value ≤0.05, and absolute value of log2 fold change ≥1. Ingenuity pathway analysis^22^ and Partek^57^ were used for analysis. Go enrichment analysis was done using Metascape web tool (https://metascape.org/) choosing GO terms with p-value <0.05. Sequencing data have been deposited in GEO under accession number GSE271995.

### CatWalk

Gait was assessed using the CatWalk XT 10.7 automated gait analysis system (Noldus Information Technology, Wageningen, The Netherlands). *HSA^CreER/+^,* β*-catenin^loxP/loxP^*mutant mice and *HSA^CreER/+^,* β*-catenin^loxP/+^* control mice received tamoxifen at P5 and were analyzed between 9-12 month of age. For each mouse, three individual assessments were conducted and for each, five runs were recorded. A successful run was determined by a duration range of 2–10 s and maximum variation of 60%. After the identification and labelling of each footprint, gait data were generated. The following parameters were analyzed for each mouse: mean speed, step sequence, stride length, print position, support and BOS (base of support).

### Statistical analyses

For quantification of immunofluorescence, all measurements were taken with confocal Z-stack projections through ImageJ 1.51. At least eight spindles from three animals of each genotype were measured. Differences were assessed by comparing animals with different genotypes with a linear mixed-effects model, with animal as a random factor. Statistical significance was defined as a p-value lower than 0.05. The analyses were done in R, v. 4.4.2, using the packages ‘lmerTest’ and ‘lme4’.

Differences in CatWalk measurements were also assessed by comparing animals with different genotypes with a linear mixed-effects model, with genotype, leg and age as fixed factors, and animal and the experimental repeats as random factors. Statistical significance was defined as a p-value lower than 0.05. The analyses were done in R, v. 4.4.2, using the packages ‘lmerTest’ and ‘lme4’.

