## Supplementary Figures for "Wnt/β-catenin is required for proprioception by coordinating the multi-compartment development of muscle spindles"

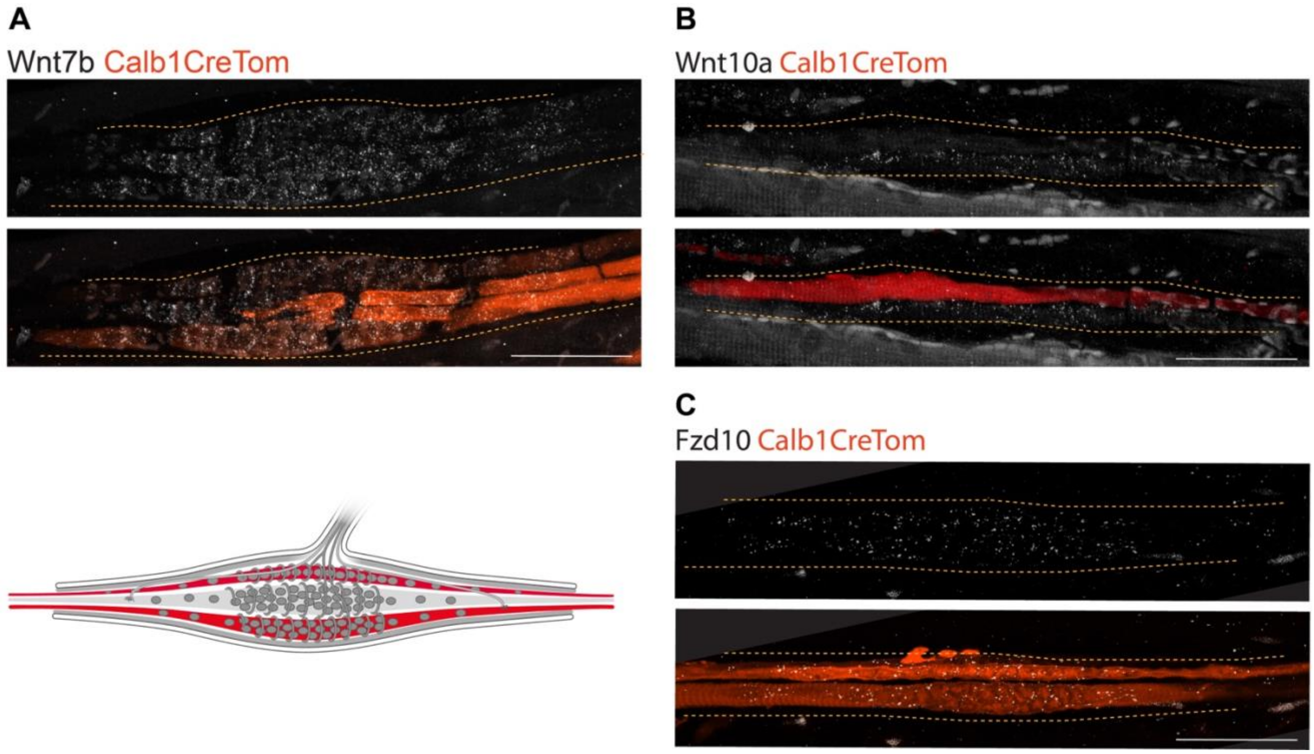


**Figure Supplement 1. Expression of Wnt genes in muscle spindles**

HCR for *Wnt7b, Wnt10a* and *Fzd10* on masseter muscle sections (A,B) and whole-mount EDL muscle (C) from *Calb1-Cre*, *Rosa26-tdTomato* mice at P25 showing expression of all three genes in intrafusal fibers. Below (A) is a schematic representation of muscle spindle with the *Calb1*$^{+}$ lineage marked in red. . Scale bar: 50 µm.

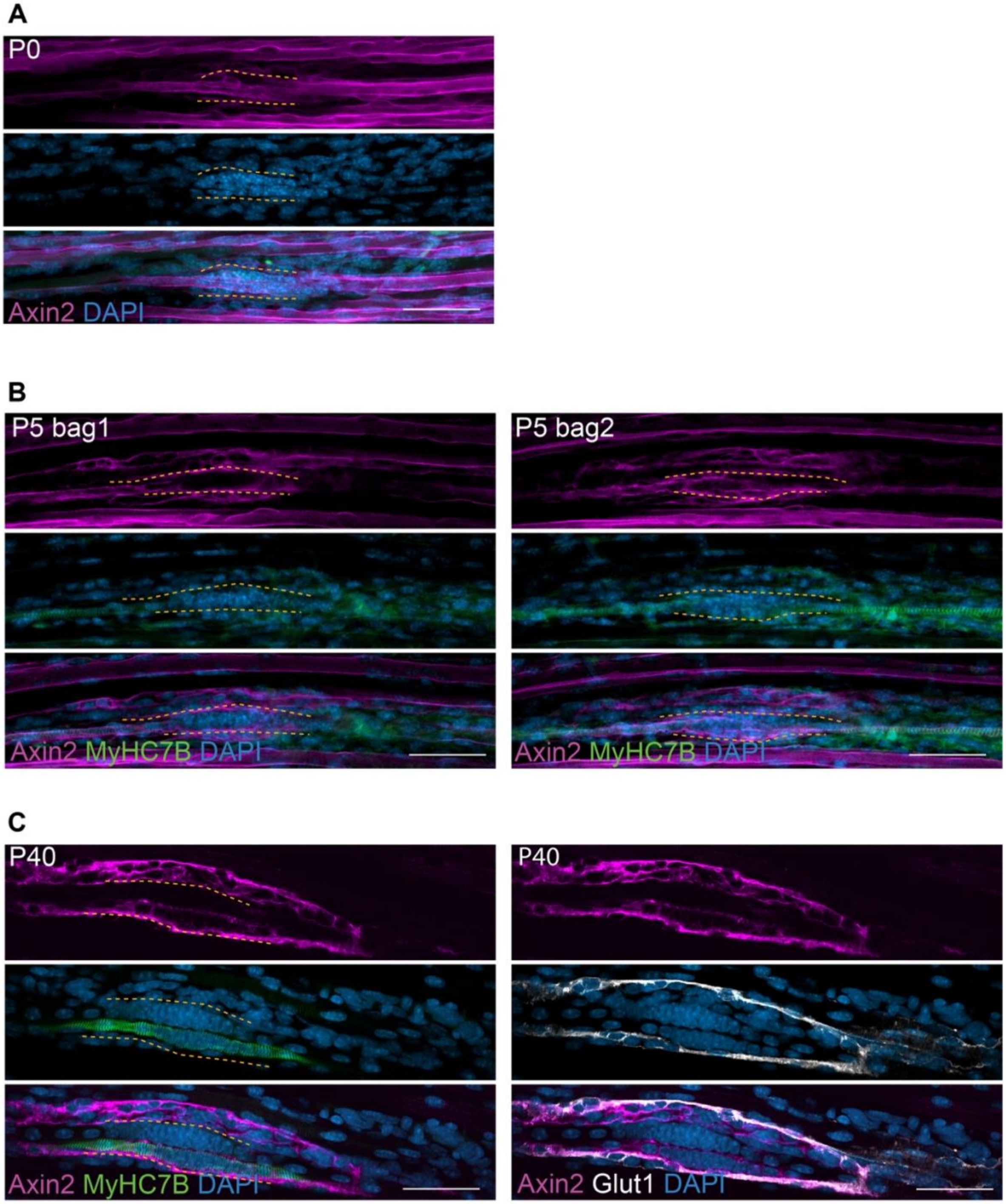


**Figure Supplement 2. Wnt/β-catenin activity in tissues of the developing muscle spindle**

Confocal images of whole-mount EDL muscles from P0, P5 and P40 *Axin2-mGFP* mice stained with antibodies against GFP (magenta) and DAPI (blue; A); GFP (magenta), MyHC7B (green) and DAPI (blue; B); or GFP (magenta), GLUT1 (white) and DAPI (blue; C). Area between dotted lines is equatorial region of intrafusal fibers. Scale bar: 50 μm.

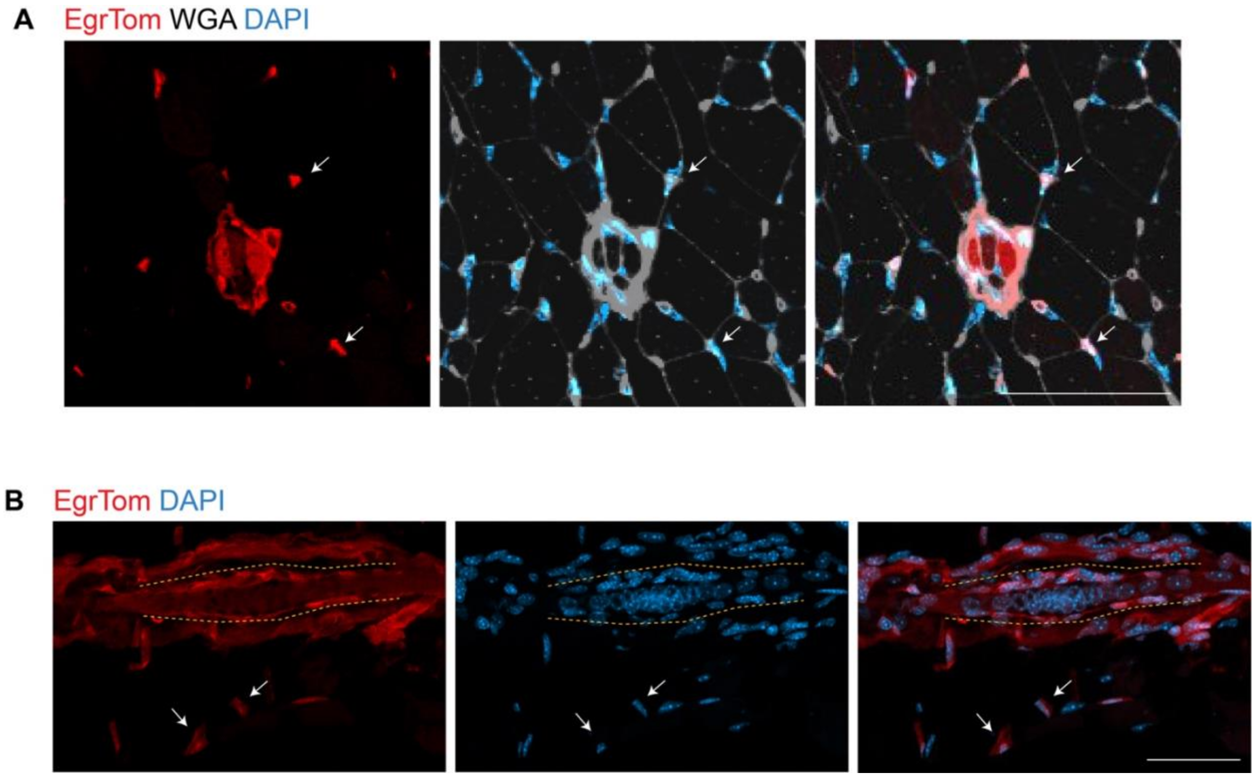


**Figure Supplement 3. Expression of *Egr3* outside the muscle spindle**

(A) Confocal images of transverse sections of forelimb muscles from P25 *Egr3-Cre, β-catenin*$^{loxP/loxP}$*, Rosa26-tdTomato* mice. Wheat Germ Agglutinin (WGA) delineates cell membrane. Arrows point at muscle interstitial cells from the *Egr3* lineage. (B) Confocal images of longitudinal sections of masseter muscles from *Egr3-Cre, Rosa26-tdTomato* mice. Dotted lines delineate muscle spindles; arrows point at *Egr3* lineage cells in the extrafusal space. Scale bar: 50 μm.

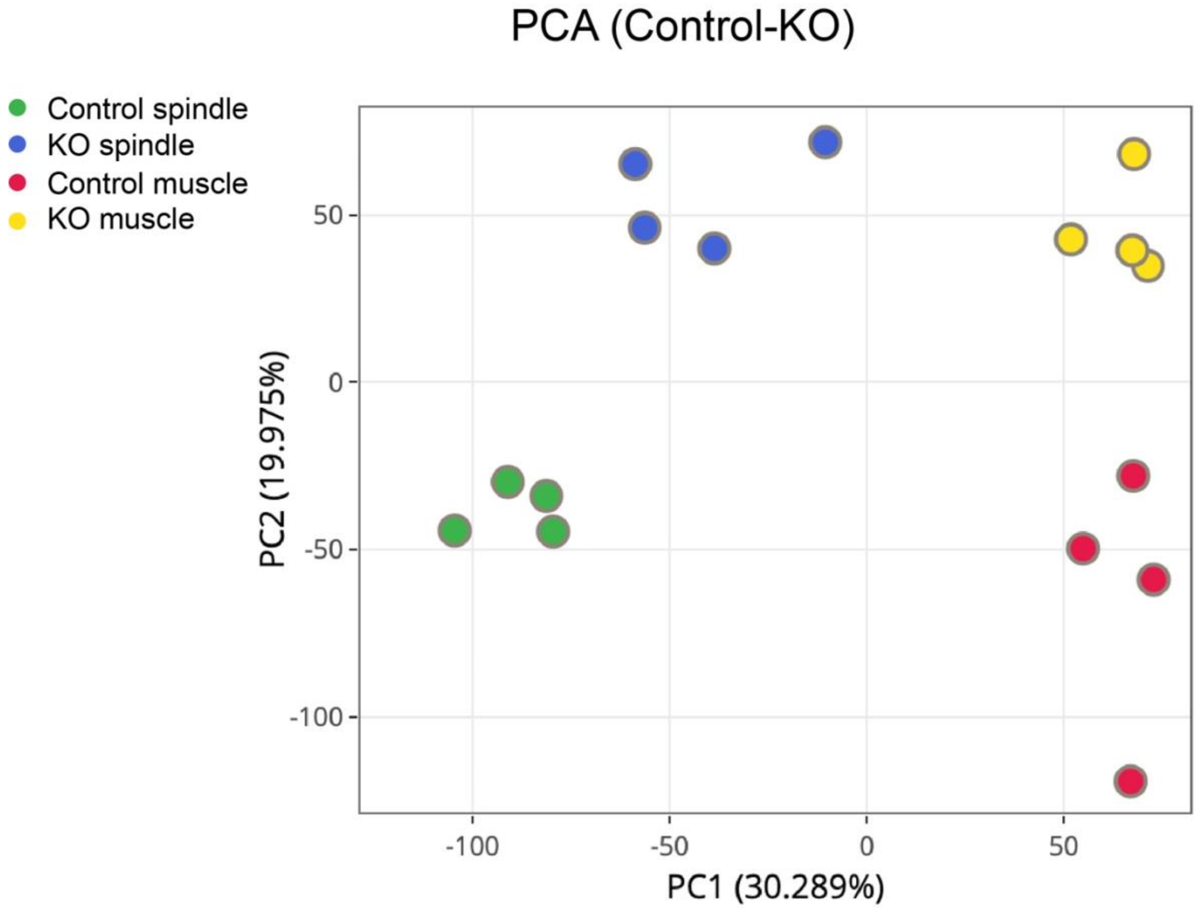


**Figure Supplement 4. Principal component analysis of bulk RNA-seq data**

Data from spindle and adjacent muscle tissue from *Egr3-Cre*, *β-catenin*$^{flox/flox}$*, Rosa26-tdTomato* mutant and control (*β-catenin*$^{flox/+}$) mice were analyzed. Green, control spindle; blue, mutant spindles; red, control muscle; yellow, mutant muscle.

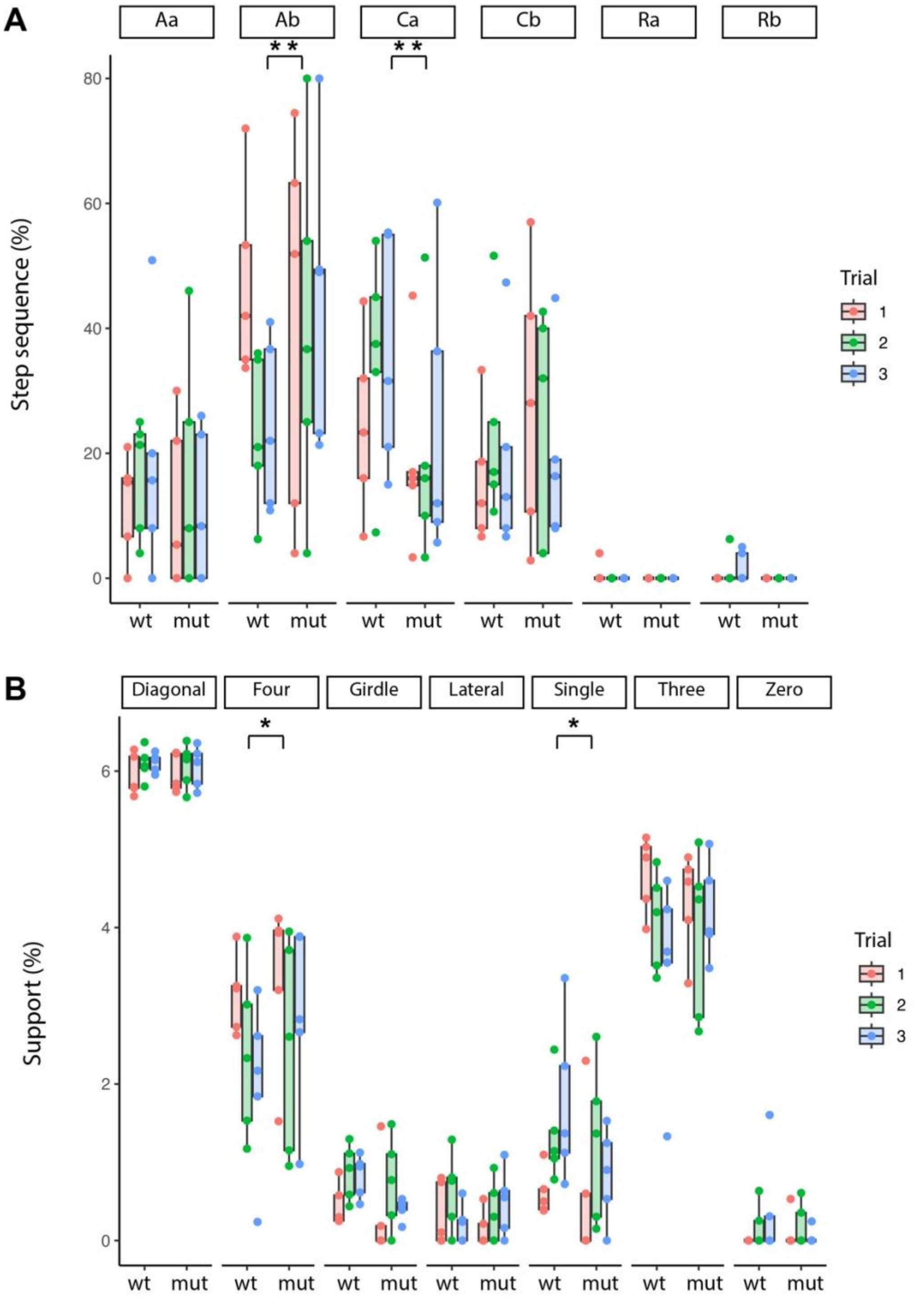


**Figure Supplement 5. Results from all individual CatWalk trials**

Each of the five animals underwent three independent proprioception assessments using the CatWalk system. Data from all trials are shown as individual points. (A) Step sequence. (B) Support pattern.
